# Lower *Vibrio* spp. abundances in *Zostera marina* leaf canopies suggest a novel ecosystem function for temperate seagrass beds

**DOI:** 10.1101/2021.03.21.436319

**Authors:** Thorsten B.H. Reusch, Philipp R. Schubert, Silke-Mareike Marten, Diana Gill, Rolf Karez, Kathrin Busch, Ute Hentschel

## Abstract

Seagrasses, a polyphyletic group of about 60 marine angiosperm species, are the foundation of diverse and functionally important marine habitats along sheltered sedimentary coasts. As a novel ecological function with high societal relevance, a role of the leaf canopy for reducing potentially harmful bacteria has recently been hypothesized. Accordingly, we tested whether or not the abundance of general bacteria and more specifically, those belonging to the genus *Vibrio* were reduced within temperate *Zostera marina* (eelgrass) meadows compared to adjacent sand flats and sampled 5 sites in the south-western Baltic Sea using SCUBA. Compared to non-vegetated area, we found an average reduction of 39% for all *Vibrio* and 63% for the potentially harmful *V. vulnificus/cholerae* subtype based on robust plate counting data on *Vibrio* selective agar. The underlying mechanism is currently elusive and clearly merits further study. Our results underline the critical importance of seagrasses in maintaining shallow water ecosystem functioning including water quality and provide further motivation for their protection and restoration.

## 1 Introduction

Marine angiosperms, or seagrasses, are the foundation of some of the most valuable coastal marine ecosystems, seagrass beds (1). As ecosystem engineers, their presence turns sedimentary bottoms with mostly infauna into a habitat featuring a rich diversity of associated animals and plants, along with substantial ecosystem services such as nursery areas, coastal protection and local carbon sequestration (2, 3). Among the many ecological services associated with seagrasses, the reduction of potentially harmful bacteria in the water column has recently been described for tropical seagrass beds (4). Lamb et al. (2017) showed that both, the abundance of possible human pathogenic bacteria as well as bacterial strains infecting marine invertebrates, were reduced in coral reef areas of the Indo-Pacific region with neighboring intact seagrass beds.

We were interested whether this effect is of general nature, and would also apply to temperate seagrasses of the northern hemisphere (5). Hence, we studied the effects of a leaf canopy of the widespread temperate seagrass *Zostera marina* (eelgrass) on water column abundance of *Vibrio* spp. bacteria, an abundant and diverse bacterial group thriving in marine waters that are generally favored by global warming and seawater freshening (6, 7). This applies particularly to our study region, the Baltic Sea, where *Vibrio* spp. abundances have already increased in the past decades due to sea surface temperature increase (6, 8), with a rate that is substantially higher than the predicted global ocean warming (9). It is therefore concerning that environmental conditions are predicted to become even more conducive to further expansion of *Vibrio* due to freshening of the brackish water body (10, 11). Alarmingly, the incidence of severe *Vibrio vulnificus* infections upon swimming or other contact with Baltic Sea waters has increased in recent years (12) which has sparked considerable media interest and potentially endangers touristic activities and revenues in the area (8).

Here, we present first pilot data on the abundance of bacterial groups belonging to the *Vibrio* spp. group in temperate seagrass beds of the south-western Baltic Sea based on a robust plate counting approach. We hypothesized that, analogous to tropical seagrass beds (4), *Zostera marina* (eelgrass) meadows would reduce the bacterial abundance in seawater within the canopy compared to adjacent sand flats. To this end, we designed a sampling program in situ by SCUBA at five locations in the southwestern Baltic Sea and assessed the abundance in seawater from inside and outside the leaf canopy at each site.

## 2 Methods

We sampled defined microhabitats associated with 5 eelgrass *Zostera marina* beds that are often interspersed with larger sandy areas in the south-western Baltic Sea. At each site, two un-vegetated and two vegetated areas were sampled in duplicates in September 2019 except for site “Gelting” where only one isolated patch of 20 m^2^ survived a recent die-off event in 2015 due to as yet unidentified causes (Table 1). Sampling points were at least 5 m away from the vegetation edge, again with the exception of the within-vegetation sampling point in Gelting that was only 2.5 m inside the remaining patch. Shoot densities in the area vary between 240 and 310 leaf shoots m^-2^, and shoot heights are 50-85 cm (13, 14).

**Table 1.** Sampling sites in the south-western Baltic Sea in September 2019.

|  |  |  | geographic coordinates |  | water depth (m) |  | environmental conditions |  |
| --- | --- | --- | --- | --- | --- | --- | --- | --- |
| site | region | date | lat (N) | long (E) | *Zostera* | sand | salinity (psu) | Water temperature (°C) |
| Gelting | Flensburg Fjord | 24 Sept | 54.7617 | 9.8789 | 0.9 | 0.6 | 22.1 | 15 |
| Kiekut | Eckenförde Bay | 4 Sept | 54.4485 | 9.8708 | 1.9-2.1 | 0.9-1.0 | 18 | 18.5 |
| Falckenstein | Kiel Fjord | 9 Sept | 54.3936 | 10.1923 | 1.1-3.3 | 0.6-0.9 | 15.3 | 18 |
| Möltenort | Kiel Fjord | 19 Sept | 54.3816 | 10.2007 | 1.1-3.1 | 0.8-1.1 | 16.6 | 15 |
| Heiligenhafen | Kiel Bight | 11 Sept | 54.3805 | 11.0077 | 0.8-3.3 | 0.5-2.3 | 14.3 | 16.8 |

Water samples were taken by SCUBA at a water depth between 0.4 and 3.4 m. Divers approached the dedicated sampling points against the prevailing current which was determined by slowly releasing seawater blobs stained with uranine (0.5 g/L) (15). Typical current speeds in the area were between 1-5 cm s^-1^. Water samples were taken by gentle suction of 100 mL water using sterile syringes 20 cm above the sediment bottom. Great care was taken that during suction sampling, SCUBA divers were well in buoyancy and did not stir up sediment. Syringes were flushed three times under water before the final sample suction.

Water samples were transferred to coolers at 4°C within 6 hrs to the laboratory where upon gentle mixing, for each sample, aliquots of 250 and 100 µL of seawater were plated onto *Vibrio* selective agar (CHROMagar_vibrio™, Chromagar Ltd. Paris, France; (16)) and onto marine broth agar, respectively. Upon a 4-day incubation at 25°C for the CHROMagar medium, and 30°C for marine broth, colonies were counted. Colonies growing on CHROMagar can unambiguously be assigned to the three major *Vibrio* spp. subtypes *V. parahaemolyticus, V. vulnificus*/*cholerae*, and *V. alginolyticus* via colony color (16).

Count data of all four response variables (3 *Vibrio* subtypes / marine broth colonies) were square-root transformed to improve variance homogeneity (17), while approximate normality was checked using residual plots. In order to accommodate the inflation of type 1-errors owing to multiple testing (18, count data were first subjected to multivariate analysis of variance (MANOVA, mean effect test) to test for an overall effect of vegetation absence /presence (hereafter “vegetation”) on bacterial & Vibrio spp. abundance. A calculation of a cross-correlation matrix explored the covariance among the 4 response variables. Subsequently, all 4 response variables were also tested in univariate analyses of variance (ANOVA) with type-III error estimation separately. At all sites, more than one subarea in one or both of the habitat categories was sampled, requiring an initial test of any “subarea(habitat)”-effect using nested analysis of variance. For all univariate response variables (colony-forming units=cfu marine broth bacteria/all Vibrio subgroups) the nested factor was not significant (all I>0.11). Based on recommendations in {Underwood, 1997 #966), we therefore pooled error terms of the nested effect and the residuals to test for the main effects “vegetation” and “site” and their interaction. Since the habitat types were partially confounded with water depth, we also tested the effect of the covariate “depth” on any of the 4 bacterial abundances in an ANCOVA model. All statistical analyses were conducted in JMP v. 10.0 (Statsoft Inc. 2012).

## 3 Results

Overall *Vibrio* spp. counts (as colony forming units, cfu 250 µl^-1^) varied between 32.3 and 66.1 among all sites, while general bacterial counts (marine broth) were substantially higher and ranged between 88.6 and 179.9 cfu 100 µl^-1^. Colonies indicative of *V. parahaemolyticus* were rare in both habitats across sites (mean±SD 1.97±2.02 cfu 250 µl^-1^), followed in abundance by the *V. vulnificus* group and the most abundant *V. alginolyticus* group (15.0±9.9 and 31.3 ±18.2 cfu 250 µl^-1^, respectively). An overall MANOVA revealed that both main factors “site” and “vegetation presence /absence” had significant effects on the overall abundance on the 4 bacterial groups tested (3 *Vibrio* subtypes / general marine bacteria on marine broth; Pillai-trace test statistic, all *P*<0.01; Supplementary Table 1). The abundance of bacteria on marine broth was only weakly correlated with overall and specific *Vibrio* group abundances (Supplementary Table S2). The abundance of the three *Vibrio* subtypes among themselves correlated only moderately with one another (all |*R*|<0.23; Supplementary Table S2).

Upon the statistically significant MANOVA, we proceeded with univariate tests on the effects of “site” and “vegetation” on the abundance of general marine bacteria and the Vibrio subgroups separately. Abundances based on colony counts on marine broth revealed a significant reduction within the vegetation (−30%; main effect vegetation, Fig. 1A, Table 2). For bacteria counted on marine broth, there was also a strong “site * vegetation presence/absence”-interaction mainly driven by the site “Kiekut” where the bacterial abundance was slightly higher within the vegetation.

**Table 2.** Results of two-way type-III ANOVAs on bacterial abundance as function of sampling site and vegetation presence /absence. df = degrees of freedom, SQ = sums of squares, MS = mean squares, *F* = *F*-ratio.

| Response variable | Effect | df | SQ | MS | *F* | *P* |
| --- | --- | --- | --- | --- | --- | --- |
| Bacteria marine broth | site | 4 | 90.31 | 22.57 | 14.13 | <.0001 |
|  | vegetation | 1 | 34.89 | 34.89 | 21.83 | <.0001 |
|  | site* veg | 4 | 47.24 | 11.81 | 8.346 | <.0001 |
|  | error | 28 | 44.75 | 1.598 |  |  |
| all *Vibrio* spp. | site | 4 | 25.74 | 6.434 | 10.83 | <.0001 |
|  | vegetation | 1 | 28.32 | 28.32 | 47.68 | <.0001 |
|  | site* veg | 4 | 19.57 | 4.892 | 8.236 | 0.0002 |
|  | error | 28 | 16.63 | 0.5940 |  |  |
| *V. parahaemolyticus* | site | 4 | 10.40 | 2.599 | 6.293 | 0.001 |
|  | vegetation | 1 | 3.680 | 3.680 | 8.902 | <.0001 |
|  | site* veg | 4 | 0.8444 | 0.2111 | 0.5112 | 0.728 |
|  | error | 28 | 11.564 | 0.4130 |  |  |
| *V. vulnificus/ cholerae* | site | 4 | 10.79 | 2.697 | 4.904 | 0.004 |
|  | vegetation | 1 | 25.38 | 25.38 | 46.15 | <.0001 |
|  | site* veg | 4 | 4.776 | 1.194 | 2.171 | 0.0983 |
|  | error | 28 | 15.40 | 0.5499 |  |  |
| *V. alginolyticus* | site | 4 | 51.57 | 12.89 | 18.54 | <.0001 |
|  | vegetation | 1 | 5.976 | 5.976 | 8.595 | 0.007 |
|  | site* veg | 4 | 14.47 | 3.618 | 5.203 | 0.0029 |
|  | error | 28 | 19.47 | 0.6953 |  |  |

**Figure 1.**
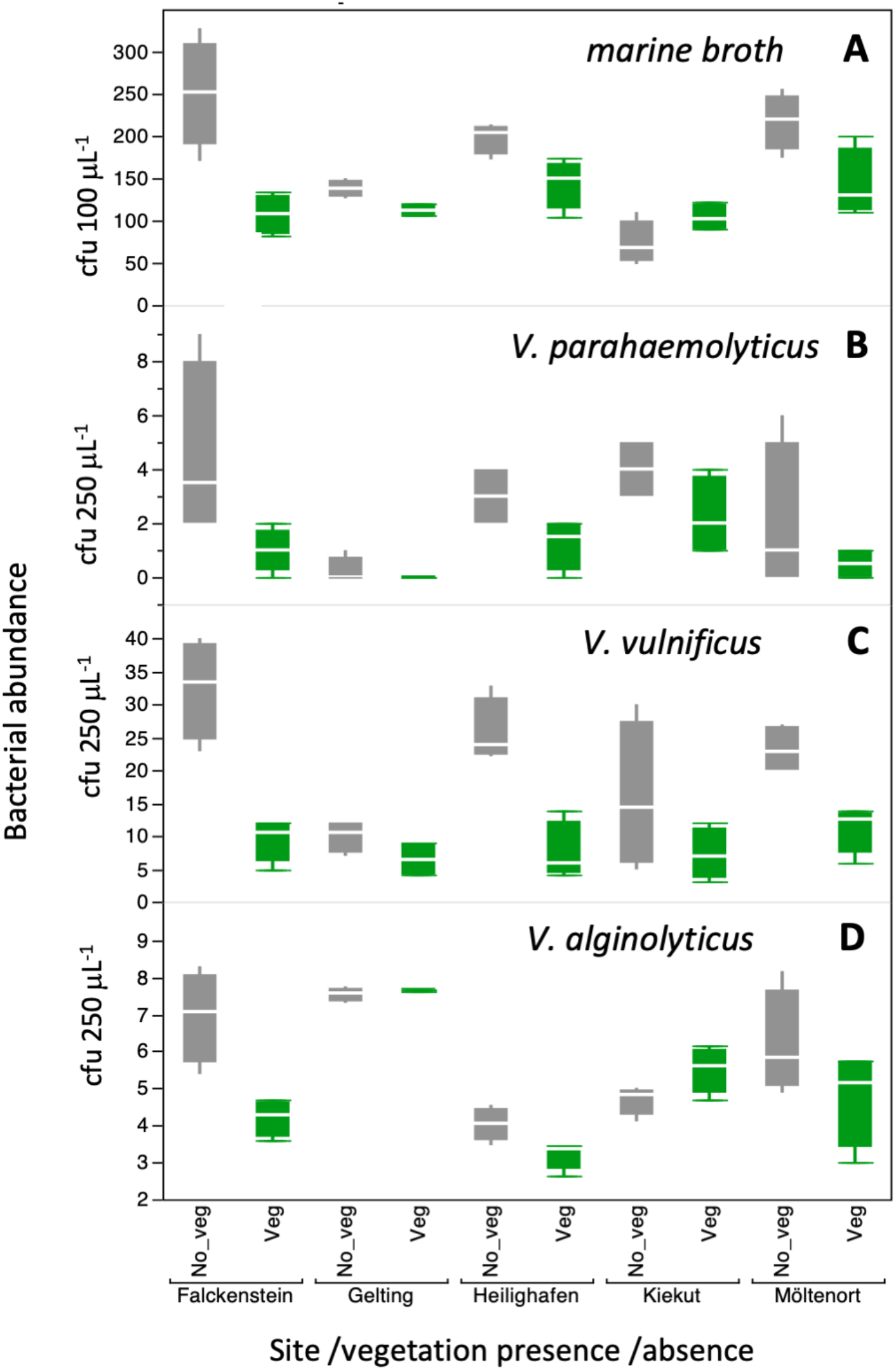
Mean abundance (±1 SD, *N*=4) of *Vibrio* spp. assessed as plate counts of colony forming units (cfu) at 5 sites in the southwestern Baltic Sea. For details on sampling sites see Table 1, for statistical analysis Table 2.

In accordance to the results of general bacterial abundance, we found that across all sites the presence of vegetation reduced the abundance of all *Vibrio* spp. by 39% compared to adjacent non-vegetated area, a reduction that was slightly higher compared to bacteria on marine broth. When broken down to subtypes, *Vibrio* spp. abundance was reduced by 63%, 61% and 23% (*V. parahaemolyticus* /*vulnificus* /*alginolyticus*, respectively), (Fig. 1B-D). While the main effect “vegetation” was significant for all *Vibrio* subtypes (all *P*_veg_ <0.007; Table 2), we also found a marked effect of “site” on all bacterial abundances, suggesting regional heterogeneities in overall bacterial abundances at the km-scale (Table 2).

The reduction in bacterial abundance was consistent across most of the sites. Specifically, despite significant overall site-differences in abundance, there was no interaction detectable among both main factors for the *V. vulnificus/cholerae* subtype (Fig. 1C). For the 2 other *Vibrio* subtypes *(V. alginolyticus* and *parahaemolyticus*) there was a significant “site*vegetation”-interaction. For *V. parahaemolyticus*, this effect originated from widely varying effect sizes among sites that varied in magnitude, while the general direction of bacterial reduction inside the leaf canopy was similar across all 5 sites (Fig. 1B). This was not the case in *V. alginolyticus*, where a closer inspection revealed that this effect was due to the site “Kiekut” where the direction of effect reversed and we observed higher abundance of bacteria within the seagrass canopy (Fig. 1D). We tested and confirmed this by excluding that site from the data set which let the statistical interaction disappear (analysis not shown).

In an ANCOVA model, we found a negative effect of water depth only on the abundance of marine broth bacteria (*r*= -0.77 ±0.26 SE) on sqrt-transformed bacterial colony counts (*P*=0.0081). In contrast, all statistical models on overall and *Vibrio* subtype abundances were not improved by including the predictor variable “water depth” into the ANCOVA (all *P*>0.13, analysis not shown).

## 4 Discussion

We found a marked reduction of all three *Vibrio* subtypes at 5 shallow water sites of the Baltic Sea during one time point in late summer as function of the presence of seagrass (*Zostera marina*) meadows. To the best of our knowledge, we provide first data that temperate seagrass beds may have the potential to reduce the abundance of potentially harmful bacteria of the *Vibrio* spp. group, consistent with earlier findings for tropical seagrass beds (4). Overall, the effects on all *Vibrio* spp. were consistent with the data on bacterial counts obtained on a general marine broth medium. An exception was the potentially harmful group of *V. vulnificus/cholerae*, where the mean reduction in cfu was with 63% the strongest in samples inside the leaf canopy compared to the abundance found in the water column above non-vegetated sandy areas outside. A similarly marked reduction was observed for *V. parahaemolyticus*, yet since the overall abundance of this *Vibrio* subtype was only ∼2 cfu 250 µl^-1^, these data have to be interpreted with caution.

Marked site effects as well as interactions of “site” with *Zostera* presence on bacterial abundance suggest heterogeneity of processes at a km-scale, hence our data should be taken as pilot data that would need confirmation in a more exhaustive spatio-temporal sampling design. Nevertheless, owing to the growing importance of *Vibrio* spp. for tourism and human health during summer months in rapidly warming northern European seas (8, 19) we consider this data set worthwhile for publication in order to motivate further studies.

Similar to the analogous role of tropical seagrass beds (4), the underlying mechanism of seagrass on surrounding water column bacteria is currently elusive. Possible non-exclusive mechanisms include (i) a simple increase of sedimentation rate through hydrodynamic attenuation of the canopy that would reduce particle load (20-22) and hence, bacteria associated with organic particles, (ii) filter feeding of fauna associated with *Z. marina* beds that removes plankton known as reservoirs for *Vibrio* or directly consumes bacteria (23, 24), or (iii) allelopathic chemicals exuding from *Z. marina* leaves (25). It is well established that the first two effects contribute to enhancement of water clarity in seagrass beds (26). The approximately similar magnitude of reduction of bacterial colonies growing on marine broth versus CHROMagar (indicative of *Vibrio* spp.) supports the hypothesis that the reduction is mainly non-selective with respect to the bacterial groups. On the other hand, for the subtype *V. vulnificus*, the reduction was even 63%, yet we currently lack the data basis to judge whether such a stronger reduction compared to general bacteria is real. In conclusion, further study is required to verify mechanisms and possible selectivity towards certain bacterial groups. Regardless of the specific mechanism, it is clear that the seagrass holobiont alters the microbiological composition in its vicinity (27), and our data are in line with these predictions.

Nature based solutions (28, 29), such as the manipulation of microbiomes (27) or the large scale restoration of macroalgae and seagrasses (28) have recently been proposed to combat various negative effects of global change and human perturbation on coastal ecosystems. Achievable goals include an improvement of water quality and the enhancement of carbon sequestration. Our study provides another example as to how the protection and possible restoration of seagrass beds (30) may help sustaining as yet understudied ecosystem services such as the suppression of bacterial load. Along with already known effects on biodiversity enhancement and improvement of water quality seagrasses may contribute to preserve the potential of coastal regions to provide tourism services without risking the health of humans visiting the ocean (19) or consuming local seafood (31).

## 6 Author Contributions

TR, PS and RK conceived the study, PS along with TR organized and conducted the sampling, DG and SMM conducted the laboratory analysis with advice from KB and UH, TR wrote the manuscript, all authors discussed the results, and commented and edited the manuscript.

## 8 Acknowledgements

Ann-Cathrin Fabricius and Vincent Fey assisted with the laboratory work, while Marlene Beer helped during field sampling. The assistance of the research divers Christian Howe, Florian Huber and Jana Ulrich in sampling acquisition is gratefully acknowledged.

## 12 Supplementary Material

Supplementary Table S1. Multivariate analysis of variance MANOVA (mean response) on bacterial abundances (marine broth /3 *Vibrio* subtypes) assessed as colony forming units. Model included “site”, “vegetation presence /absence” and their interaction as fixed effects.

Supplementary Table S2. MANOVA: matrix of partial correlation coefficients among 4 response variables marine broth bacteria /*Vibrio* spp. subtypes.

Supplementary Table S3. Original data set of colony forming units on marine broth and on Vibrio-selective agar (CHROMagar_vibrio).

## 13 Data Availability Statement

Original data are available in Supplementary Table S3 and have also been uploaded onto PANGAEA (doi: xx).

